# Co-clinical FDG-PET Radiomic Signature in Predicting Response to Neoadjuvant Chemotherapy in Triple Negative Breast Cancer

**DOI:** 10.1101/2021.06.11.448077

**Authors:** Sudipta Roy, Timothy D. Whitehead, Shunqiang Li, Foluso O. Ademuyiwa, Richard L. Wahl, Farrokh Dehdashti, Kooresh I. Shoghi

## Abstract

**Purpose:** We sought to exploit the heterogeneity afforded by patient-derived tumor xenografts (PDX) to optimize robust radiomic features associated with response to therapy in the context of a co-clinical trial and implement PDX-optimized image features in the corresponding clinical study to predict and assess response to therapy using machine-learning (ML) algorithms.

**Methods:** TNBC patients and subtype-matched PDX were recruited into a co-clinical FDG-PET imaging study to predict response to therapy. One hundred thirty-one imaging features were extracted from PDX and human segmented tumors. Robust image features were identified based on reproducibility, cross-correlation, and volume independence. A rank importance of predictors using ReliefF was used to identify predictive radiomic features in the preclinical PDX trial in conjunction with ML algorithms: classification and regression tree (CART), Naïve Bayes (NB), and support vector machines (SVM). The top four PDX-optimized image features, defined as radiomic signatures (RadSig), from each task were then used to predict or assess response to therapy. Performance of RadSig in predicting/assessing response was compared to SUV_mean_, SUV_max_, and lean body mass normalized SUL_peak_ measures.

**Results:** Sixty-four out of 131 preclinical imaging features were identified as robust. NB-RadSig performed highest in predicting and assessing response to therapy in the preclinical PDX trial. In the clinical study, the performance of SVM-RadSig and NB-RadSig to predict and assess response was practically identical and superior to SUV_mean_, SUV_max_, and SUL_peak_, measures.

**Conclusions:** We optimized robust FDG-PET radiomic signatures (RadSig) to predict and assess response to therapy in a context of a co-clinical imaging trial.

**DECLARATIONS:** *Funding:* This work was supported by NCI grants U24CA209837, U24CA253531, and U54CA224083; U2CCA233303, and K12CA167540; Siteman Cancer Center (SCC) Support Grant P30CA091842; and Internal funds provided by Mallinckrodt Institute of Radiology.

*Conflicts of interest/Competing interests.:* None.

*Availability of data and material:* All the co-clinical data will be available for download through the Washington University School of Medicine Co-Clinical Imaging Research Resource web portal at https://c2ir2.wustl.edu/, co-clinical database (CCDB).

*Code availability:* Not applicable.

*Authors’ contributions:* Conceptualization: SR, FOA, KIS; Methodology: SR, TDW, SL, KIS; Formal analysis and investigation: SR, KIS; Writing - original draft preparation: SR; Writing - review and editing: RLW, FD, KIS; Funding acquisition: RWL, FOA, SL, KIS; Resources: SL; Supervision: FD, KIS. All authors read and approved the final manuscript.

*Ethics approval:* All studies were performed with approval from the Washington University Humans subjects research committee and animal studies committee.

*Consent to participate:* Informed consent to participate in the study was obtained from all participants.

*Consent for publication:* Not applicable.

## INTRODUCTION

Triple Negative Breast Cancer (TNBC) is a highly heterogeneous and aggressive cancer characterized by poor outcome and higher relapse rates compared to other subtypes of breast cancer. Pathological complete response (pCR) is often used as a critical endpoint in the treatment of TNBC following NAC as it is often associated with favorable long-term outcome. Therefore, it is critical to identify patients who will respond to NAC therapy to avoid the use of ineffective treatments. Intratumoral heterogeneity is regarded as a major factor in tumor progression and resistance to NAC [1]. Towards that end, advanced quantitative imaging (QI) strategies, including extraction of image features, or radiomics, have been employed to characterize tumor heterogeneity and to predict/assess response to therapy [2, 3].

We designed a co-clinical trial to assess the efficacy of docetaxel/carboplatin therapy in patients with TNBC and patient-derived tumor xenografts (PDX) generated from TNBC patient biopsies. Co-clinical trials are an emerging area of investigation in which a clinical trial is coupled with a corresponding preclinical trial to inform the corresponding clinical trial [4–10]. The emergence of PDXs as co-clinical platforms is largely motivated by the realization that established cell-lines do not recapitulate the heterogeneity of human tumors and the diversity of tumor phenotypes [11]. Indeed, numerous investigations have demonstrated that PDX accurately reflect patients’ tumors in terms of the histomorphology, gene expression profiles, and gene copy number alterations [12–16], as well as ability to predict therapeutic response in patients, especially when a clinically relevant drug dosage is used [17–19]. To that end, the National Cancer Institute’s (NCI) Patient-Derived Models Repository (https://pdmr.cancer.gov), EuroPDX (https://www.europdx.eu), academic institutions, and numerous commercial entities have launched wide-ranging PDX and repositories to advance the use of PDX in oncologic applications.

One of the objectives of the co-clinical trial, which is still underway, is to predict response to therapy using [^18^F]fluorodeoxyglucose (FDG) with positron emission tomography (PET). We previously identified six TNBC subtypes including 2 basal-like (BL1 and BL2), an immunomodulatory (IM), a mesenchymal (M), a mesenchymal stem–like (MSL), and a luminal androgen receptor (LAR) subtype through molecular signatures of TNBC subtypes [20]. The use of PDX in preclinical imaging offers numerous advantages in translational imaging research, chief among them is retention of human tumor heterogeneity [12, 16, 21], which can be exploited to develop image metrics of response to therapy. Thus, the primary objective of this work was to utilize PDX to optimize robust radiomic features of tumor heterogeneity indicative of response to therapy in preclinical PDX trials.

The scheme outlined in Figure 1 highlights the paradigm we undertook in this effort. We used the co-clinical imaging trial to define, for the first-time, parallels in radiomic features between preclinical and clinical imaging. To address the primary objective, we characterized the reproducibility, cross correlation (auto-correlation), and volume dependency of FDG-PET radiomic features in PDX. Optimal radiomic features were then used in ML algorithms to define radiomic signatures (RadSig) of response to therapy in the preclinical PDX trial. With the RadSig at hand, we performed an interim analysis to implement radiomic signatures optimized in the preclinical PDX trial to predict response to therapy in the clinical arm. Our findings suggest that RadSig performed significantly better than standardized uptake value (SUV) measures to predict (using baseline metrics) and assess (difference in image metrics) response to therapy.

**Figure 1:**
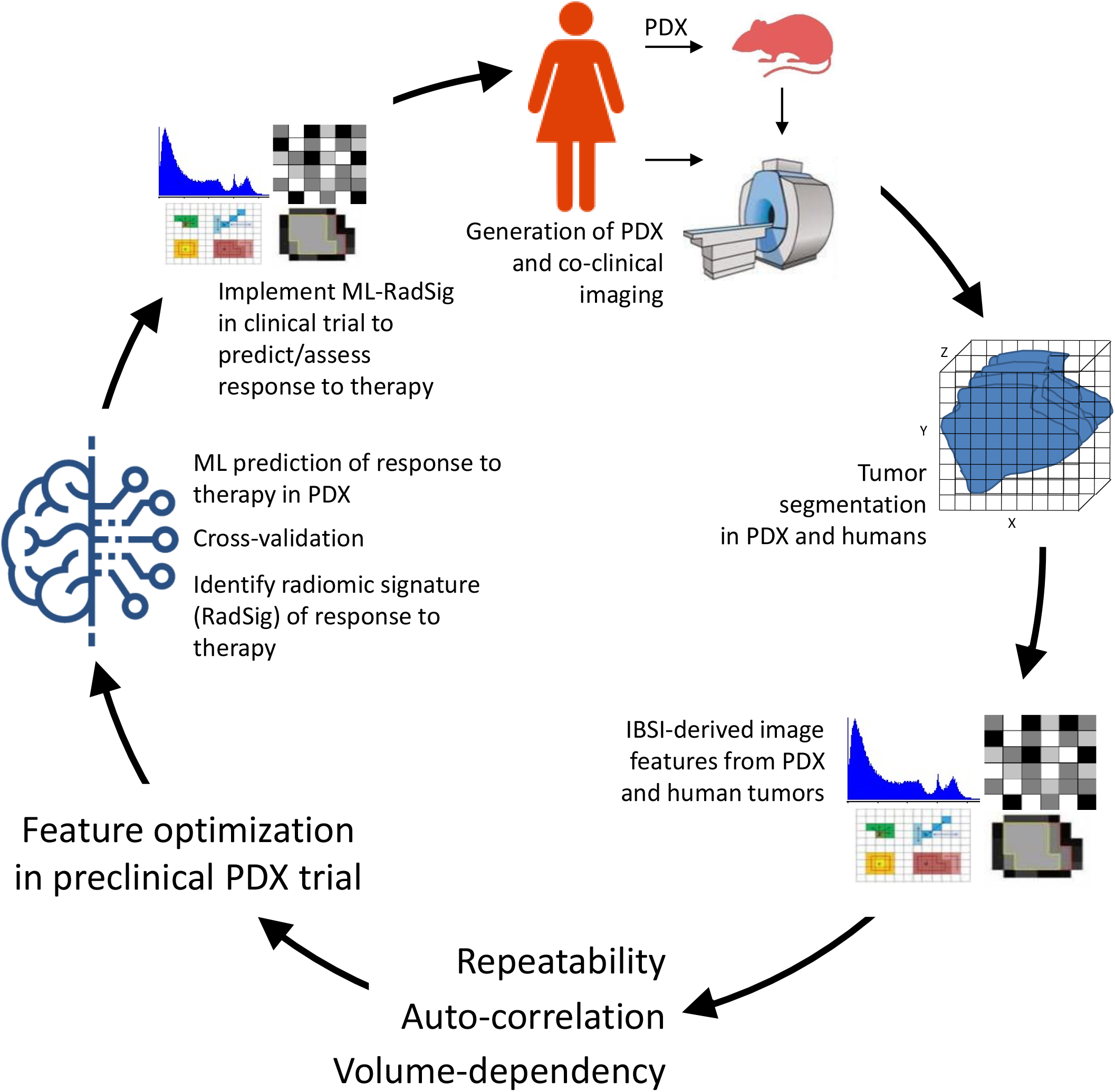
Overview of methodology in optimizing radiomic features in the co-clinical trial. TNBC PDX were generated from human tumor biopsies. Tumor were segmented following co-clinical imaging to extract radiomic features. Radiomic features were extracted per IBSI guidelines. Repeatability, cross-correlation, and volume-dependency were performed to identify the robust features. ReliefF, and then ML were used to predict/assess the response to therapy in PDX and to identify radiomic signature (RadSig). RadSig was implemented in the clinical trial to predict/assess response to therapy.

## METHODS

### Co-clinical protocol

The co-clinical design is outlined in the scheme of Figure 2A and described below. TNBC PDX subtypes were identified as described previously [22] based on molecular signature analysis. TNBC subtypes include: basal-like (BL1 and BL2), an immunomodulatory (IM), a mesenchymal (M), and a luminal androgen receptor (LAR) subtype.

**Figure 2:**
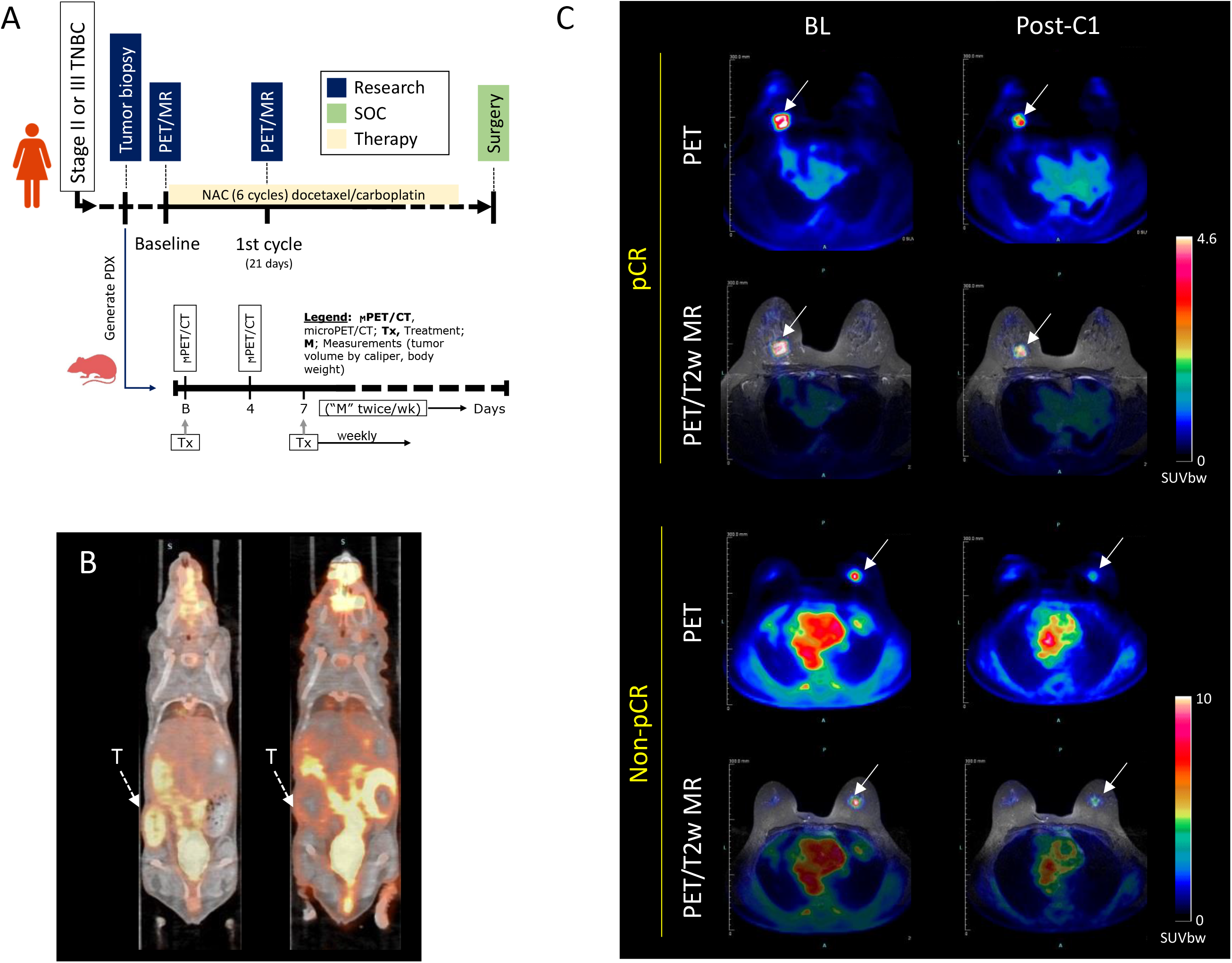
Co-clinical imaging. (A) Co-clinical imaging study design. Stage II or III TNBC patients were recruited into the study for 6 cycles of docetaxel/carboplatin therapy. Imaging timepoints are indicated on the timeline. PDX are generated from patient tumor biopsies to assess response to therapy with imaging at baseline and +4 days post-therapy. (B) representative preclinical PET/CT and (C) clinical PET/MR images of response to therapy. Tumor is indicated by white arrow.

#### Preclinical imaging

Small animal PET/CT was performed on the Inveon microPET/CT scanner as described previously [22]. Briefly, four hours prior to imaging session, food was removed from metabolism cages while water was given ad libitum. Mice were anesthetized with 2-2.5% isoflurane by inhalation via an induction chamber. Anesthesia was maintained throughout the imaging session by delivering 1%–1.5% isoflurane via a custom-designed nose cone. A heat lamp was used to maintain body temperature. TNBC PDX were injected with ^18^FDG (6.66 – 8.14 MBq) by tail vein immediately before a 0-60 min dynamic small animal PET acquisition. Images were reconstructed with a 3D OSEM algorithm with a Ramp filter at 0.5 cutoff and voxel size of ~0.08mm. In therapeutic studies, TNBC PDX (N=29) were imaged at baseline (BL) and four days (4D) following start of therapy (Figure 2A). Docetaxel (20mg/kg IP)/carboplatin (50mg/kg IP) was administered following BL imaging and weekly for a period of four weeks. Tumor volumes were measured bi-weekly. All animal experiments were conducted in compliance with the Guidelines for the Care and Use of Research Animals established by Washington University’s Animal Studies Committee.

#### Clinical imaging

Twenty stage II or III TNBC patients were recruited into an ongoing co-clinical trial (ClinicalTrial.gov ID # NCT02124902). A secondary goal of the co-clinical trial was to assess the performance of FDG-PET in predicting/assess response to therapy. Simultaneous FDG-PET and MR imaging protocols were implemented on the Siemens Biograph mMR. Subjects were imaged at baseline (BL) prior to therapy and between the first cycle (C1) and second cycle of docetaxel/carboplatin for a total of 6 cycles (21 days per cycle). At each imaging time point, patients fasted for ~4hrs prior to injection of ~10mCi of FDG. After an uptake period, patients were positioned prone on the PET/MR scanner. FDG-PET imaging was performed starting at 30min to 70min post FDG administration. Default Dixon sequence was used for attenuation correction. Images were reconstructed to produce four 10min frames. In parallel with FDG-PET acquisition, T1-weighted (T1w) and T2-weighted (T2w) MR acquisitions were performed. The co-clinical trial is ongoing; however, we performed an interim analysis to assess the performance of PDX-optimized FDG-PET image features to predict/assess response to therapy in the clinical arm. Quantification of clinical MR data is not included at this stage since preclinical imaging did not include MR acquisition.

### Image analysis and extraction of radiomic features

#### Preclinical imaging

Static 10min PET/CT images obtained 50min post-administration of FDG (representative image in Figure 2B) were processed in two-steps. In the first step, co-registered PET/CT images were analyzed using the Inveon Research Workplace (IRW) software (Siemens Healthcare). Volumes of interest (VOIs) were manually drawn on co-registered PET/CT images to include tumor(s). Second, VOIs and individual voxels were normalized to SUV in MATLAB using the relation: SUV = [activity (Bq / mL)] × [animal weight (g)] / [injected dose (Bq)].

#### Clinical imaging

Tumor VOIs were manually drawn on 20min static PET images obtained by averaging two 10min frames 50-70min post-administration of FDG (representative image in Figure 2C). To ensure harmonization of preclinical and clinical pipelines, IRW was used to segment tumors on PET/MR images. Mean SUV (SUV_mean_) for the entire tumor was calculated as per above. Peak SUV was normalized to lean body mass (SUL_peak_) based on positron emission tomography response criteria in solid tumors (PERCIST) [23].

#### Extraction of imaging features

One hundred thirty-one imaging features were extracted from preclinical and clinical tumors. These include: one hundred twenty radiomic features, tumor volume, metabolic tumor volume, and nine SUV metrics as tabulated in Supplemental Table S1. Radiomic features were determined per the image biomarker standardization initiative (IBSI) guidelines [24, 25]. Equal-probability quantization algorithms to quantize raw data into gray level (Ng) were implemented using histeq MATLAB function. Resampling to isotropic voxel size in all three directions was applied to all higher order features. Thirty-seven first order features were extracted directly from raw data. All higher order features were extracted after applying fixed quantization of gray level Ng=64.

### Robustness of radiomic features

We evaluated the robustness of radiomic features in terms of reproducibility (test-retest), cross-correlation, and the dependency on tumor volume. Robust radiomic features were then used as predictors of response to therapy.

#### Test-retest

A preclinical test-retest protocol was implemented to optimize the reproducibility of radiomics features. PDX (N=40) were imaged on consecutive days (Day 1 and Day 2) in identical conditions.

#### Cross-correlation

The cross-correlation between features was determined using Spearman correlation. A threshold Spearman correlation of ρ≥0.9 and significance value P<0.001 were chosen as significantly high correlation between features.

#### Volume-dependent radiomic features

Radiomic features were regressed against their corresponding tumor volumes. Linear or nonlinear functional forms were used to fit all significant volume-dependent features.

### Prediction and assessment of response to therapy

#### Prediction vs. assessment of response to therapy

We make a distinction between predicting and assessing response to therapy. In predicting response to therapy, BL imaging features were used to predict response to therapy in either the preclinical or the clinical arm. In assessing response to therapy, the change (Δ) in imaging feature between on-treatment (4D in preclinical and post C1 in clinical) and BL was used to predict response to therapy in either the preclinical or the clinical arm.

#### Classification of response to therapy

In preclinical studies, end-point caliper volume change from start of treatment was considered as surrogate of response to therapy with response to therapy corresponding to >20% decrease in volume; partial response corresponding to ≤|20|% change in volume, and no response corresponding to >20% increase in volume. Baseline radiomic features and change in radiomic features between 4D post-treatment and baseline scans were used as the predictive criterion for ML algorithms. In clinical studies, pCR was used to determine response to therapy.

#### Feature selection

In preclinical studies, the Relief-based algorithm (RBA) [26] was used to select a subset of features as inputs to the ML algorithms. A relevance threshold (τ=0.05) [27] was used to select most relevant weighted features to facilitate in expansive modeling, reduce overfitting, and make the task tractable for inputs in ML algorithms. These optimal features were used to predict response or assess response to therapy using BL and difference between on-treatment and BL optimal features, respectively.

#### Machine learning for outcome prediction

The ML algorithms used in this study include CART [28], SVM [29], and NB [30]. In CART, Gini index was used at each partition to determine splitting criteria with a binary threshold of CART. In implementing SVM, radial Basis Function (RBF) kernel was used to make the hyperplane decision boundary between the classes. Objective function L2-norm regularization was used to overcome overfitting problem. CART, SVM, and NB work well with datasets as low as N=20 [31]. Ten-fold cross-validation was used to avoid overfitting the ML model [32].

### Statistical Analyses

#### Robustness of features

Lin’s concordance correlation coefficient (LCC) [33] was used to assess reproducibility using Stata version 12.1. LCC≥0.7 was considered as a threshold of reproducible radiomic feature [34, 35]. As indicated above, cross-correlation between features was evaluated using the Spearman correlation ρ≥0.9 at significance value P<0.001. To display clusters of correlations, hierarchical clustering of the Spearman correlation heatmap was performed. In evaluating volume-dependency of features, the Akaike Information Criterion (AIC) and Bayesian Information Criterion (BIC) were calculated for each functional form, and the appropriate model was selected based on the minimum value of AIC and BIC. The Spearman correlation (ρ) was used to determine the correlation between each feature and tumor volume.

#### Performance metrics of response to therapy prediction

Common performance metrics including accuracy, F-score, sensitivity, specificity, precision, and negative predictive value (NPV) were used to assess performance of response to therapy [20]. The performance of the radiomic features was additionally compared with SUV_mean_, SUV_max_, and SUL_peak_ based on PERCIST [23].

## RESULTS

### Reproducibility of preclinical radiomic features

Test-retest was performed to assess the reproducibility of radiomic features using LCC as a measure of reproducibility. Ninety-four out of 129 radiomic features (72.9%) were identified as reproducible with LCC≥0.7. The frequency of correlations along with the cumulative percent is displayed in Figure 3A. Approximately 22% of features were highly reproducible with LCC≥0.9. The reproducibility by class of features is depicted in Figure 3B. Figures 3C depicts the LCC values of all reproducible radiomic features. Supplemental Table S1 summarizes the reproducibility of all 131 features.

**Figure 3:**
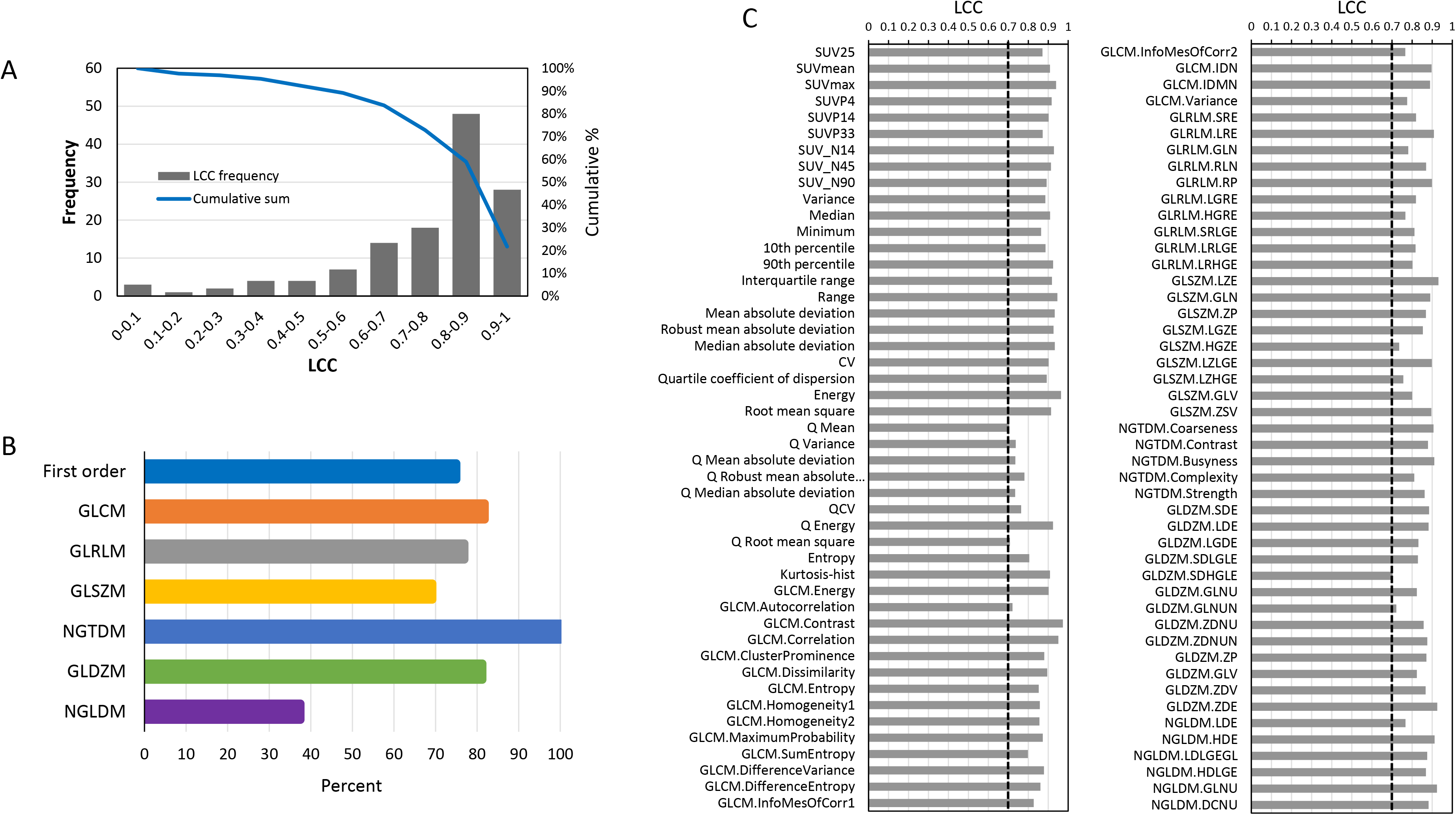
Repeatability of preclinical radiomic features. (A) Frequency (bar plot, left Y-axis) of repeatability by LCC values and cumulative percent (solid line, right Y-axis). (B) Percent of repeatable features with LCC≥0.7 by class (left Y-axis labels). (C) Reproducible radiomic features with LCC≥0.7 (denoted by dashed vertical black line).

### Cross-correlation between features (preclinical and clinical)

We ascertained the cross-correlation between 129 features using the Spearman correlation (ρ). Highly correlated features (ρ≥0.9) were removed and reduced to 94 features from 129 features. Hierarchical clustering of the Spearman correlation heatmap is shown Figure 4. Twenty-one clusters were identified in the preclinical heatmap (Figure 4A) and similarly 21 clusters were identified in the clinical cross-correlation heatmap (Figure 4B). Membership of features to clusters is available in Supplemental Table S2. The distribution of Spearman correlations is available in Figure 4C and 4D for preclinical and clinical cross-correlations, respectively.

**Figure 4:**
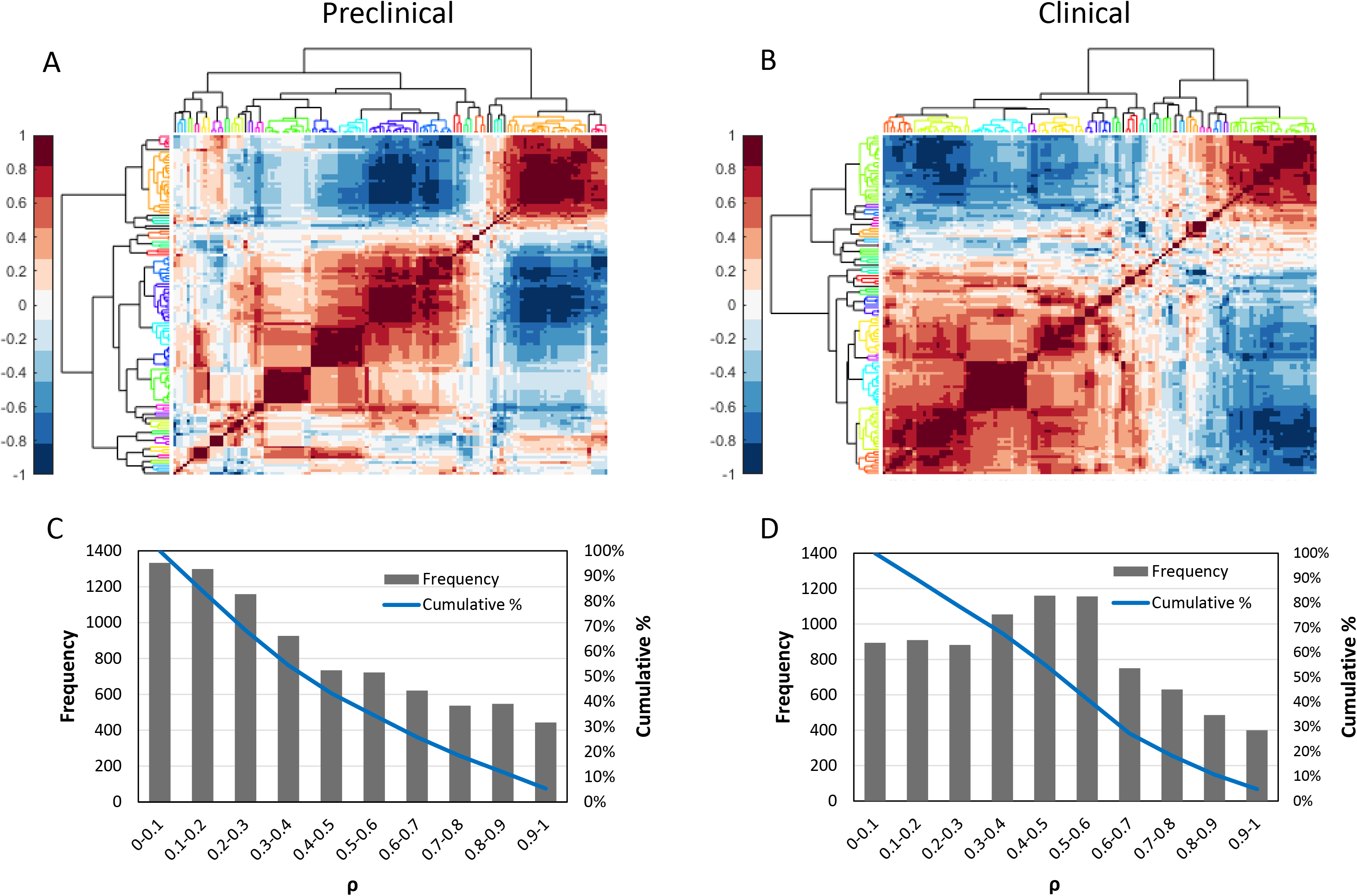
Cross-correlation between radiomic features. Hierarchical clustering of the Spearman cross-correlation heatmap for (A) preclinical and (B) clinical. The dendrogram shows the clustering, and each color represents a different cluster. The frequency and cumulative sum at each Spearman correlation (ρ) is displayed in (C) for preclinical and (D) for clinical.

### Volume-dependent radiomic features (preclinical and clinical)

In total, 10 radiomic features were highly correlated to volume (ρ>0.9; P<0.001). The functional form of the volume dependency and corresponding goodness-of-fit measures for preclinical and corresponding clinical images is shown in Figure 5, which was similar for both preclinical and clinical features. Supplemental Table S3 summarizes the statistical analyses for the correlations.

**Figure 5:**
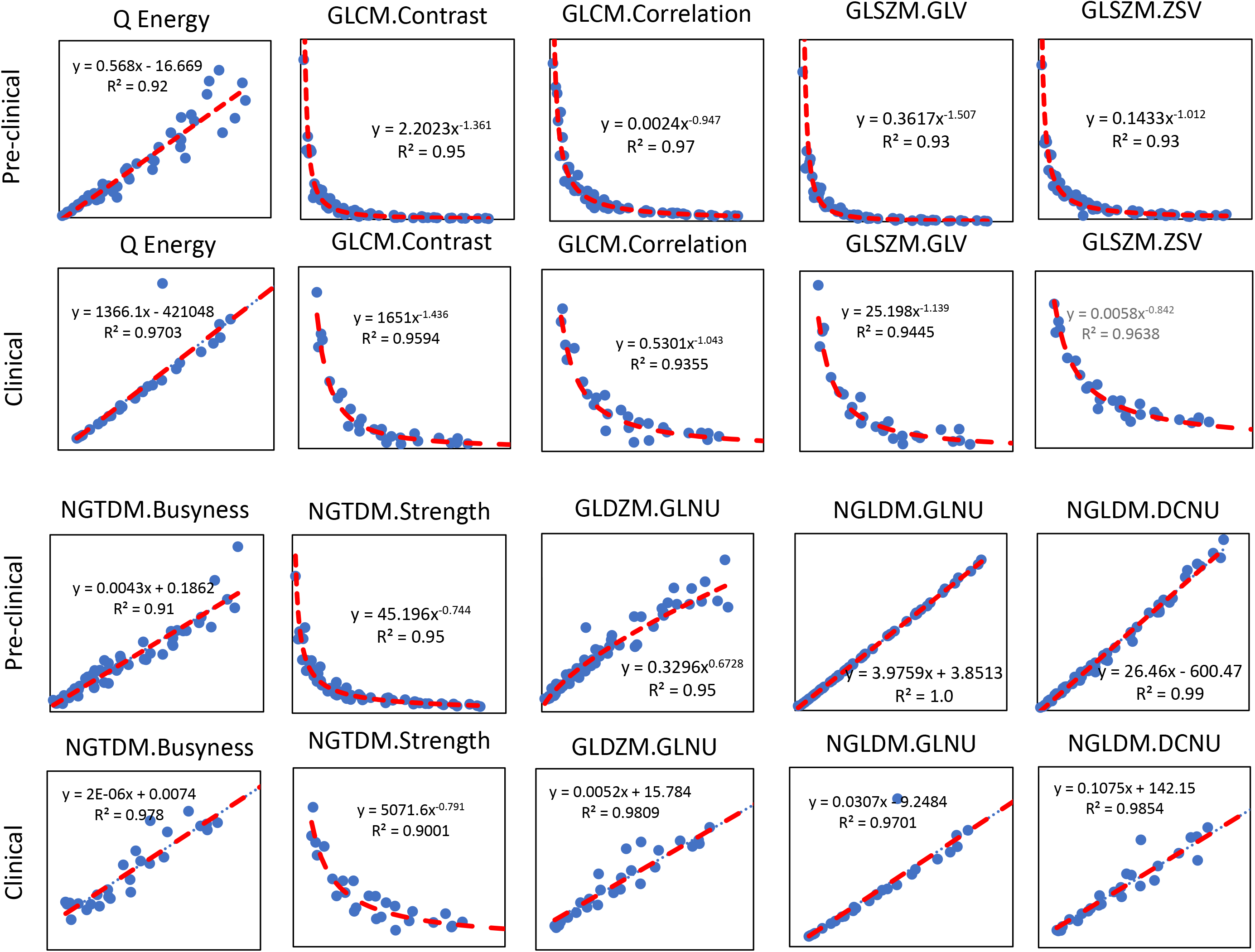
Volume-dependent preclinical and clinical radiomic features. The correlation between features and tumor volume was assessed for preclinical and clinical segmented tumors. Ten common preclinical and clinical features exhibited high correlation (ρ≥0.9) with tumor volume.

### Prediction and assessment of response to therapy

At the intersection of robustness analyses, 62 of the 129 (48.06%) features were found to be optimal and were passed to ReliefF feature selection followed by ML. ReliefF rank importance identified top performing 15 features for prediction (based on BL features) and assessment (based on 4D-BL features) of response to therapy (Figures 6B and 6C, respectively). The rank importance of radiomics features is given in Supplemental Table S4.

**Figure 6.**
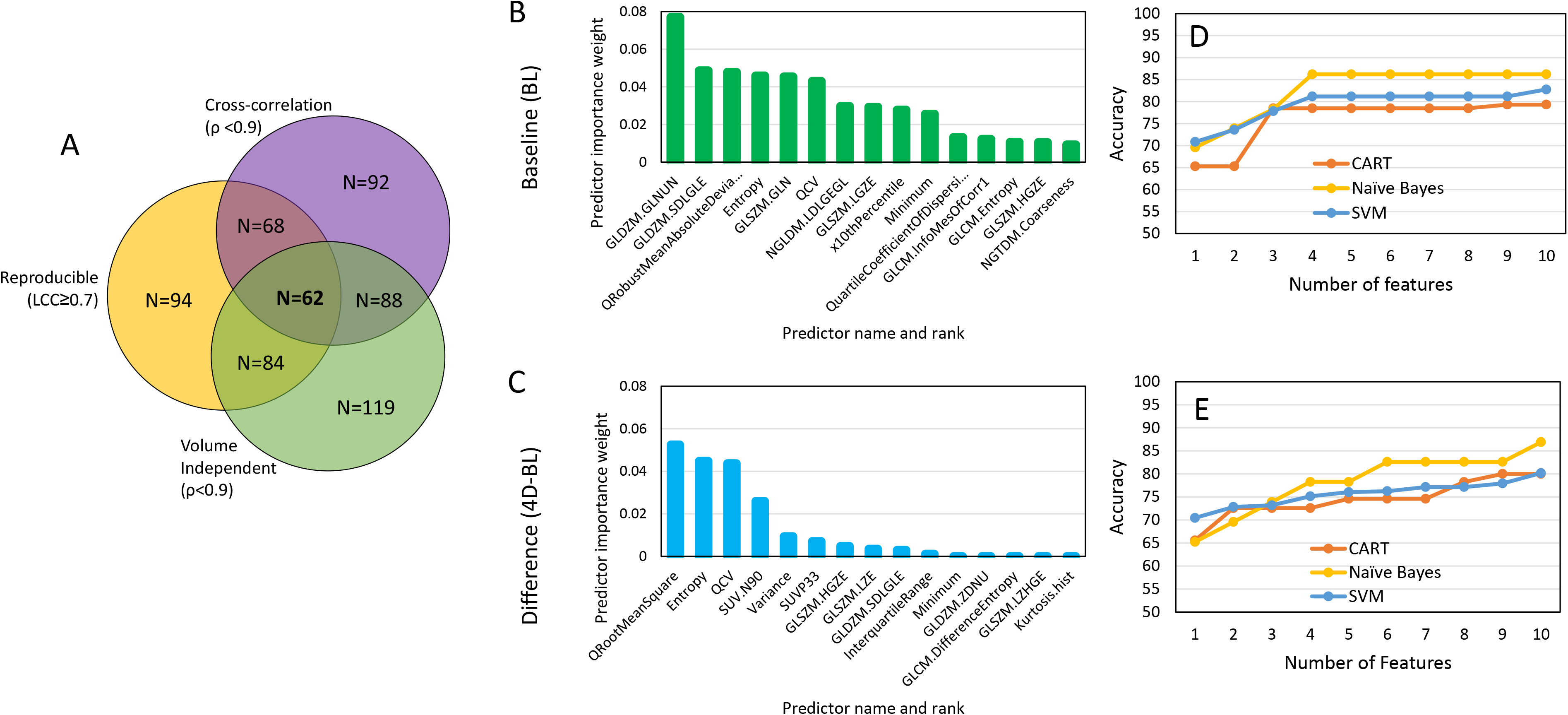
ML-based selection of radiomic features. (A) At the thresholds defined within to screen for volume-dependency, repeatability, and cross-correlation, we identified 62 optimal features. Implementation of relief-based algorithm (RBA) to identify a subset of features as inputs to ML-based prediction (B) and assessment (C) of response to therapy. Accuracy of ML algorithms CART, SVM, and Naïve Bayes to predict (D) or assess (E) response to therapy as a function of number of radiomic features in PDX.

#### Preclinical PDX studies

The accuracy of ML in predicting/assessing response to therapy as a function of the number of radiomic features is depicted in Figures 6D and 6E for BL and 4D-BL, respectively. The number of radiomic features to maximize prediction accuracy saturated at 4 features (Figure 6D) with NB exhibiting the highest accuracy at 86.21 %, followed by SVM, and CART. In contrast, the accuracy of assessing response to therapy (4D-BL) increased with increasing number of radiomic features; the accuracy of NB is 86.9% followed by SVM and CART (Figure 6E). We opted to compare performance between prediction and assessment (i.e., BL versus 4D-BL) using the least number of robust features. For this reason, Table 1 tabulates the performance of the ML algorithms to predict/assess response prior to and following optimization for robust features using only the top 4 radiomic features for each classification (prediction versus assessment of response). The set of 4 radiomic features from each task (prediction and assessment of response) make up the Radiomic Signature (RadSig). As tabulated in Table 1, RadSig performs as well as, or marginally better than, non-optimized features (all features) in predicting response. The performance of prediction/assessment of response to therapy stratified by TNBC subtype is tabulated in Supplemental Table S5 and highlights differences in prediction by TNBC subtype.

**Table 1:** Accuracy of predicting (BL) and assessing (4D-BL) response to therapy using top 4 radiomic features.

| Methods | All features |  | RadSig |  |
| --- | --- | --- | --- | --- |
|  | Prediction | Assessment | Prediction | Assessment |
| CART | 80.34 | 74.86 | 78.48 | 72.57 |
| Naïve Bayes | 82.62 | 82.76 | 86.21 | 78.26 |
| SVM | 78.48 | 78.45 | 81.14 | 75.13 |

The performance of RadSig in comparison to SUV_mean_, SUV_max_, and SUL_peak_ for the top two performing ML algorithms (NB and SVM) is summarized in Figure 7. NB performed marginally better than SVM in predicting/assessing response to therapy (Figure 7A) in the preclinical PDX trial. The percent increase in predicting/assessing response to therapy relative to SUV_mean_, SUV_max_, and SUL_peak_ is depicted in Figure 7B for NB. NB-RadSIG improved prediction of response by over 60% in all performance measures. In assessing response to therapy, RadSig performed better than SUV_mean_ in most performance criteria and marginally better than SUL_peak_ and SUV_max_ (Figure 7B). Thus, RadSig has greater impact in predicting response to therapy than assessing response to therapy. Full performance data is available is Supplemental Table S6. We then performed an interim analysis of the ongoing clinical trial to implement PDX-optimized RadSig to predict/assess response to therapy using ML.

**Figure 7:**
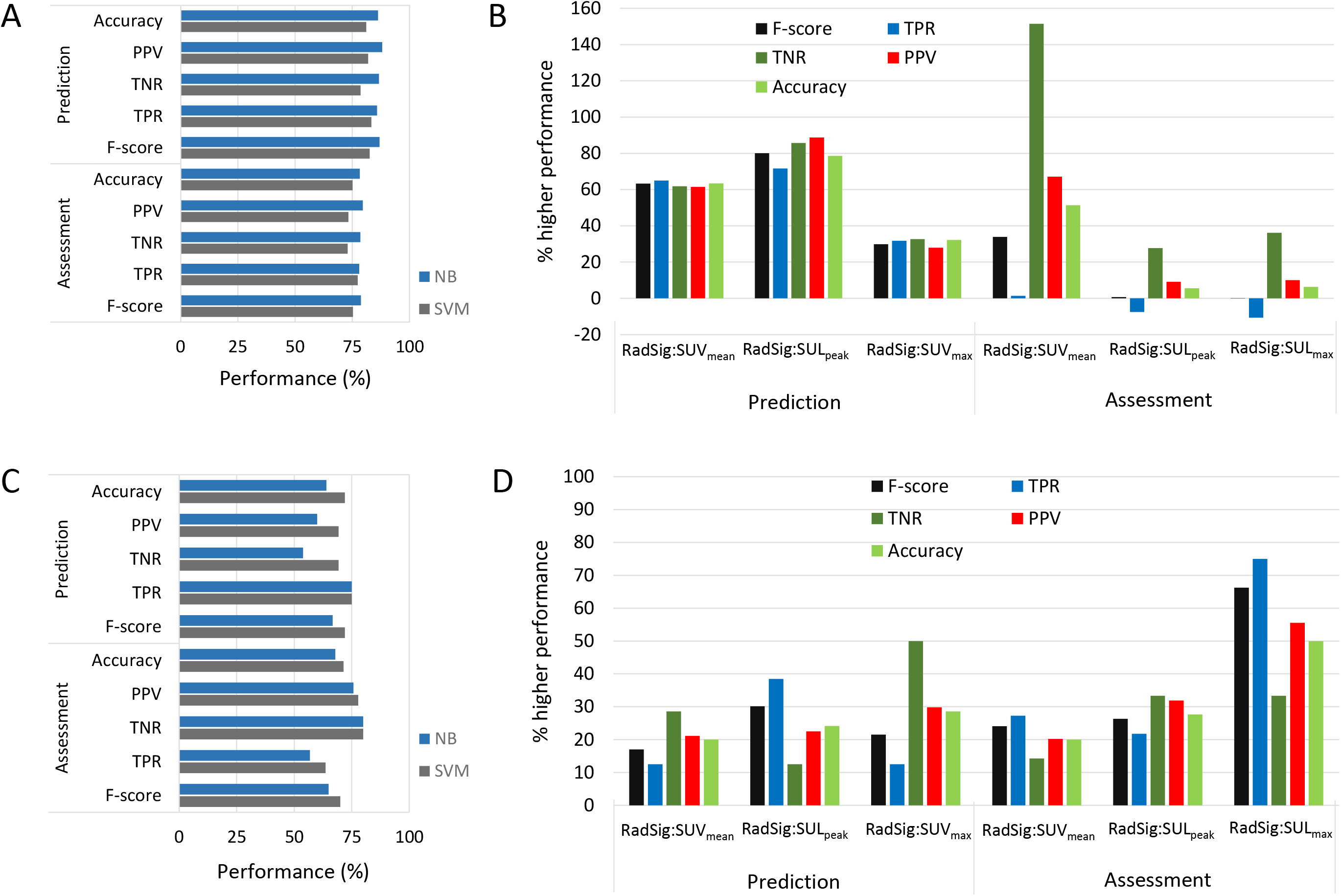
Performance of ML algorithms. (A) Performance of RadSig in predicting/assessing response to therapy in the preclinical PDX trial with NB and SVM. (B) Percent improvement in NB-RadSig prediction/assessment of response relative to SUV_max_, SUV_mean_ and SUL_peak_. (C) similar to (A) but for the clinical investigation. (D) similar to (B) but using SVM.

Table 2 summarizes patient characteristics, pathological response, SUV metrics at BL, and percent change in SUV metrics between on-treatment (post C1) and BL for the interim analyses. Of the twenty patients, ten patients exhibited pCR; however, all patients exhibited reduction in SUV. Average percent (±1SD) reduction in the non-pCR group was −46.94±21.56, −53.20±19.91, and −51.33±19.78 for SUV_mean_, SUL_peak_, and SUV_max_, respectively; and −57.70±14.83, −60.32±16.47, and −66.16±13.74 for SUV_mean_, SUL_peak_, and SUV_max_, respectively. Figure 7 also depicts the performance of the ML algorithms in predicting and assessing response to therapy in the clinical arm (Figure 7C). The performance of SVM and NB with RadSig as a predictor were marginally similar, although overall SVM performed better than NB when using SUV metrics as predictors (Supplemental Table S6). SVM-RadSig exhibited higher prediction rates of response to therapy relative to SUV_max_, SUV_mean_ and SUL_peak_ in all performance measures (20-40% higher), as well as in assessing response to therapy (15-75% higher) (Figure 7D). Overall, RadSig performed better than SUV metrics in predicting and assessing response to therapy.

**Table 2:** Patient characteristics, pathologic response, and SUV metrics

| Stage at Diagnosis | Grade at Diag. | pCR | BL $SUV_{mean}$ | BL $SUL_p$ | BL $SUV_{max}$ | %$\Delta$ $SUV_{mean}$ | %$\Delta$ $SUL_{peak}$ | %$\Delta$ $SUV_{max}$ |
| --- | --- | --- | --- | --- | --- | --- | --- | --- |
| IIB (T2N1) | 2 | No | 1.86 | 1.57 | 4.03 | NA | NA | NA |
| IIB (T2N1) | 3 | Yes | 6.59 | 4.58 | 11.27 | -60.97 | -68.11 | -57.82 |
| IIIA (T3N1) | 3 | No | 10.55 | 12.34 | 26.41 | NA | NA | NA |
| IIA (T2N0) | ? | Yes | 8.77 | 7.80 | 21.48 | -75.49 | -82.21 | -86.77 |
| IIB (T2N1) | 3 | No | 3.18 | 1.64 | 5.31 | -4.32 | -13.19 | -9.40 |
| IIB (T2N1) | 3 | No | 2.92 | 1.29 | 4.54 | -38.67 | -41.17 | -41.77 |
|  |  |  | 8.01 | 6.43 | 15.55 | -40.82 | -44.37 | -42.90 |
| IIB (T2N1) | 3 | No | 8.58 | 6.20 | 17.02 | -64.92 | -70.33 | -73.40 |
|  |  |  | 8.91 | 6.62 | 20.11 | -62.72 | -68.17 | -61.72 |
| IIA (T2N0) | 3 | Yes | 2.49 | 1.61 | 5.25 | NA | NA | NA |
| IIA (T2N0) | 3 | Yes | 4.17 | 3.06 | 12.45 | -48.84 | -48.53 | -64.23 |
| IIA (T2N0) | 3 | Yes | 3.89 | 3.12 | 8.62 | -57.76 | -59.91 | -73.72 |
| IIB (T2N1) | 3 | No | 2.85 | 2.59 | 6.28 | -41.45 | -55.68 | -55.97 |
|  |  |  | 2.48 | 2.03 | 5.03 | -27.04 | -37.81 | -49.88 |
| IIA (T2N0) | 3 | No | 8.66 | 7.56 | 21.87 | -75.82 | -82.48 | -81.35 |
| IIB (T2N1) | 3 | Yes | 10.29 | 8.32 | 28.33 | -74.14 | -76.39 | -81.07 |
|  |  |  | 3.64 | 2.02 | 5.65 | -58.93 | -57.11 | -55.17 |
|  |  |  | 8.00 | 5.60 | 16.06 | -71.75 | -77.22 | -77.14 |
| IIA (T2N0) | 3 | Yes | 1.86 | 1.65 | 4.18 | -30.42 | -36.91 | -53.02 |
| IIA (T2N0) | 3 | No | 9.80 | 7.74 | 17.69 | -67.65 | -64.44 | -54.15 |
| IIA (T2N0) | 3 | Yes | 7.19 | 5.21 | 14.57 | -46.36 | -42.49 | -43.23 |
| IIA (T2N0) | 3 | No | 7.98 | 7.26 | 18.46 | -45.99 | -54.32 | -42.74 |
| IIA (T2N0) | 3 | Yes | 2.91 | 2.03 | 5.24 | -40.23 | -40.04 | -58.66 |
| IIB (T2N1) | 3 | Yes | 6.19 | 5.53 | 11.28 | -69.81 | -74.57 | -76.91 |
| IIB (T2N1) | 3 | No | 2.44 | 1.99 | 5.53 | NA | NA | NA |

## DISCUSSION

The emergence of co-clinical models is largely motivated by the realization that established cell-lines do not recapitulate the heterogeneity of human tumors and the diversity of tumor phenotypes [11] and that better oncology models are needed to support high-impact translational cancer research [12, 16, 21]. An underlying premise in the co-clinical study design is that the heterogeneity of the human tumor is retained in PDX. Indeed, tumor genomic and pathological investigations have confirmed that PDX recapitulate the heterogeneity of human tumors [12–16] and that these can be used to a better inform cancer biology, therapeutic design [17–19], and therefore by extension imaging studies, albeit with some limitations [21]. With that in mind, in this this work, we exploited the heterogeneity of TNBC PDX subtypes to 1) identify robust radiomic features in preclinical TNBC PDX; 2) optimize RadSig-ML algorithms to predict response to therapy in PDX; and 3) implement PDX-optimized RadSig to predict/assess response to therapy in the corresponding clinical trial.

To our knowledge, this study represents the first such effort to optimize radiomic features in preclinical PET imaging to predict/assess response to therapy in TNBC PDX. We recently characterized the dependency of preclinical MR radiomic features on tumor volume [36]. In this work, we confirmed dependency of preclinical PET radiomic features on tumor volume with strikingly similar clinical parallels. This is particularly relevant in longitudinal studies during which tumor volumes will change with the course of the disease or following therapy. Ideally volume-independent features should be used as to not bias image features longitudinally. We further evaluated the cross-correlation of preclinical and clinical radiomic features with the goal of reducing the dimensionality of features. Finally, we evaluated the repeatability of radiomic features in preclinical PET imaging to identify robust features for inclusion in ML-based prediction of response to therapy. At the thresholds defined within to screen for volume-dependency, repeatability, and cross-correlation, we identified 62 optimal features to predict/assess response to therapy.

RBF [26] was used to rank image features using three ML algorithms as to their relevance in predicting/assessing response to therapy. Our data suggests that overall SVM performed better than NB and CART in predicting response to therapy. We used the top four ML-RBF-optimized radiomic features—referred to as radiomic signature (RadSig)—from each task (prediction vs. assessment) to either predict or assess response to therapy. RadSig performed significantly better in predicting response to therapy in the preclinical and clinical arm, as well as in assessing response to therapy in the clinical arm. Antunovic et al. [37] reported the utility of FDG-PET radiomic features to assess response to therapy using four different models in 79 patients with heterogenous breast cancer subtypes. The reported area-under the curve of an ROC analysis ranged from 0.70 to 0.73. Li et al. [38] recently assessed the utility of both PET and CT radiomic features to predict response to therapy in a retrospective study that included 100 heterogenous breast cancer patients. The PET/CT radiomic predictors achieved a prediction accuracy of 87% on the training split set and 77% on the independent validation set.

In the small, albeit homogenous, dataset of TNBC patients where PDX-optimized radiomic features were implemented in the clinical imaging arm, we observed an impressive accuracy of 72% and 71% when predicting and assessing response, respectively, compared to SUV metrics. We were unable to perform a validation test on an independent dataset; however, the primary objective was to compare the performance of all predictive metrics in the training phase. In addition, we did not report MR radiomic features in this work. However, prediction of response to therapy can be further enhanced through integration of MR imaging features [39], liquid biopsies such as circulating tumor DNA (ctDNA) analyses [40], and molecular/genomic features of tumors [41], all of which are an active area of investigation. Finally, numerous recent studies have documented that pCR rates varied with breast cancer molecular subtypes. TNBC and HER2-positive molecular subtypes have shown to have higher pCR rates after NAC [42]. Importantly, several studies have demonstrated an association between imaging features and molecular phenotypes, risk of recurrence, and prognosis [43–45]. Interestingly, our PDX studies similarly suggest that response to therapy (and prediction thereof) is a function of the TNBC subtype, however further studies are needed to support this hypothesis and the utility of radiomic features in classifying TNBC subtypes. With that in mind, one of the most critical aspects in practical implementation of radiomics is a consensus on the most effective features and their standardization.

## CONCLUSIONS

We identified robust FDG-PET radiomic features from an ongoing co-clinical (PDX and human) trial to predict and assess response to therapy. The number of radiomic features to maximize accuracy was further optimized in the preclinical PDX trial to yield ML radiomic signatures (RadSig) of response to therapy. We then implemented RadSig in an interim analysis of the corresponding clinical trial. The performance of SVM-RadSig in predicting/assessing response to therapy was superior to SUV_max_, SUV_mean_ and SUL_peak_ metrics in the clinical setting; however, given the small sample size additional studies are warranted to further validate RadSig and potentially integrate with multi-scale features to enhance prediction/assessment of response to therapy.

## Acknowledgements

The authors acknowledge the staff of the Preclinical Imaging Facility and the Center for Clinical Imaging Research (CCIR) at Mallinckrodt Institute of Radiology (MIR), Washington University School of Medicine for performing imaging studies.

---

**Supplemental Table S1:**
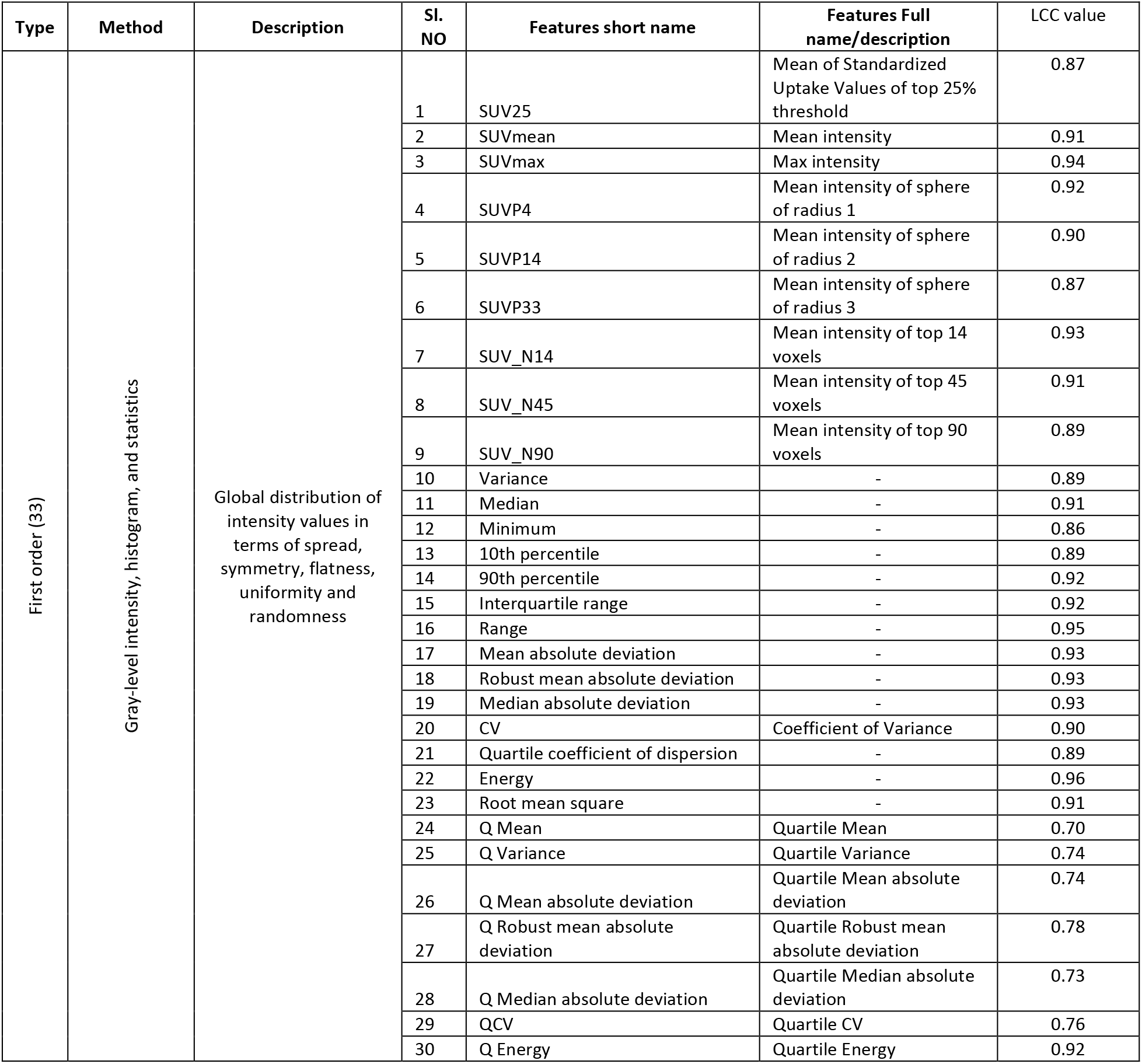
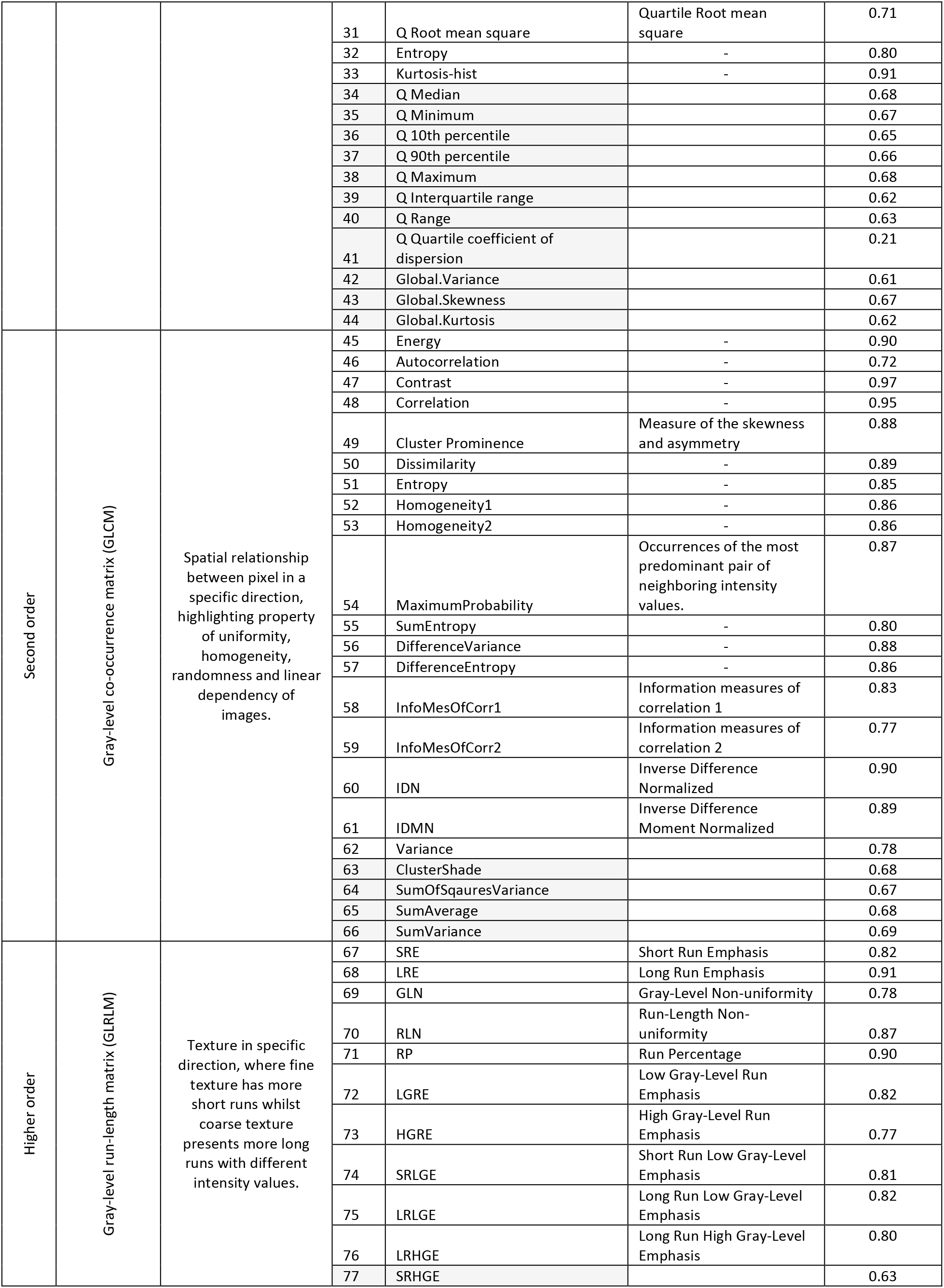
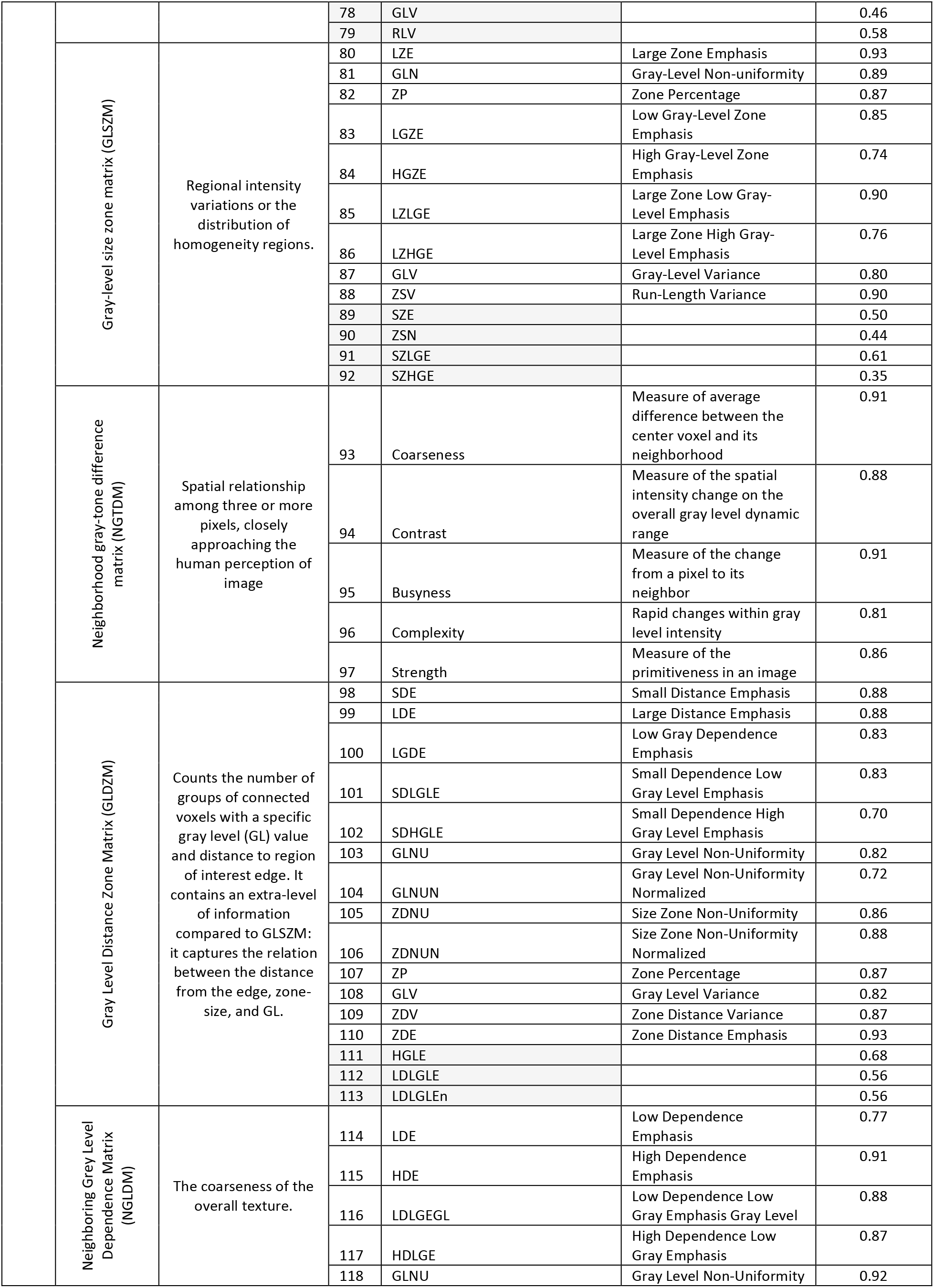
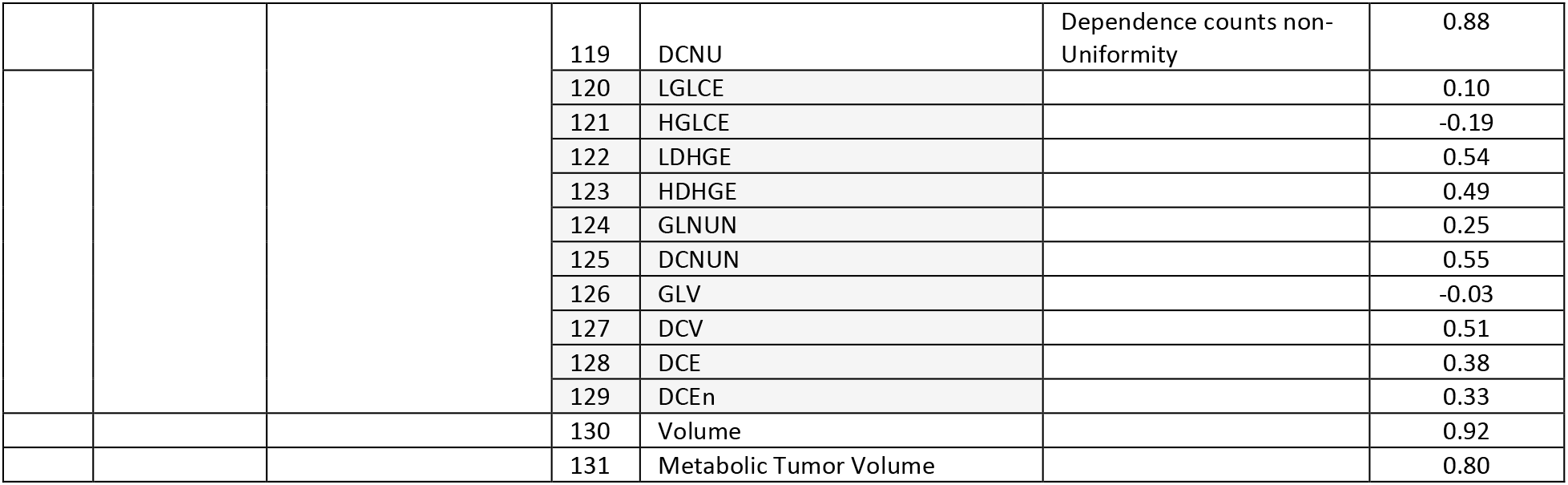
Radiomic features. Light shaded color radiomics were non reproducible features.

**Supplemental Table S2:**
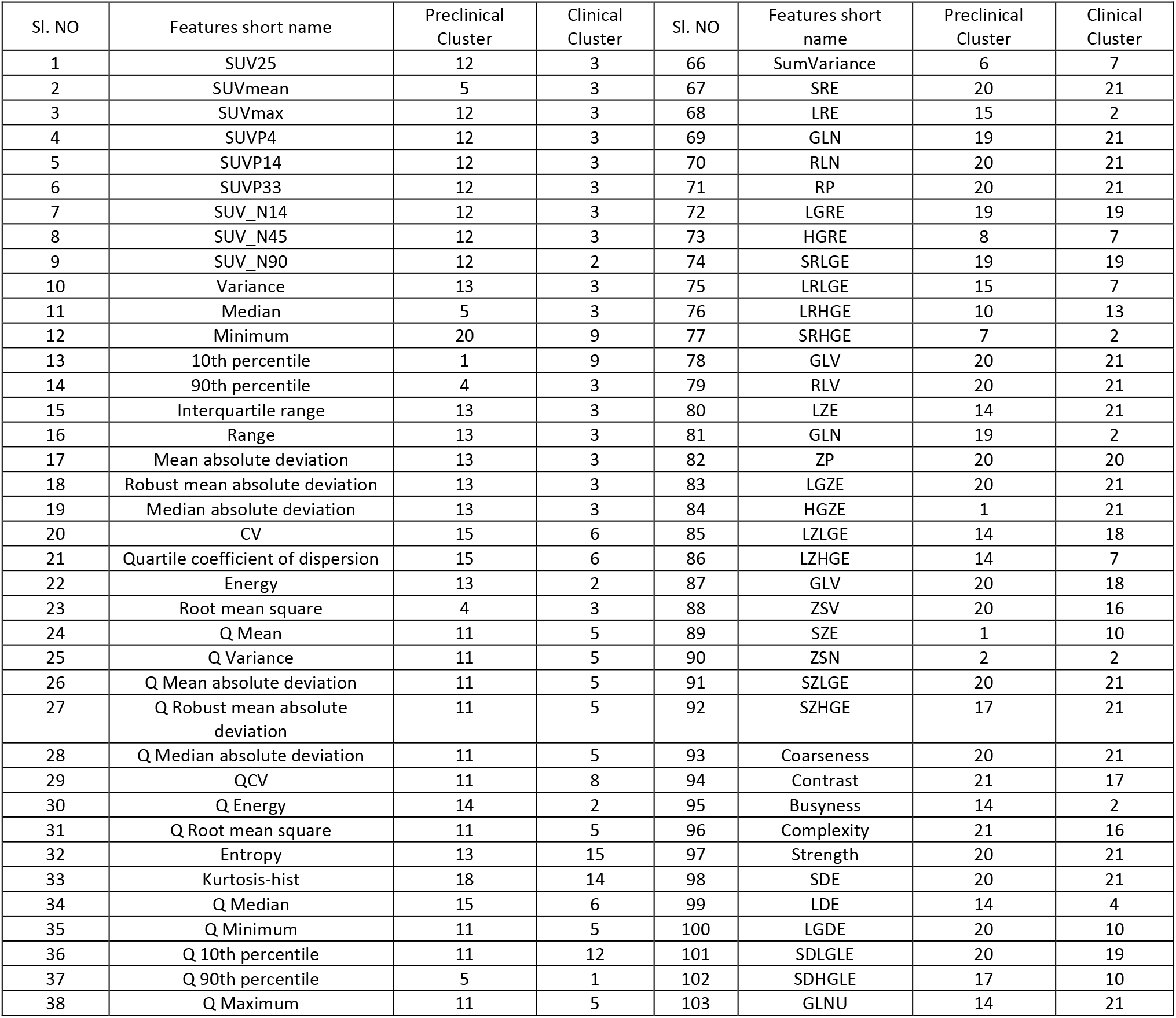
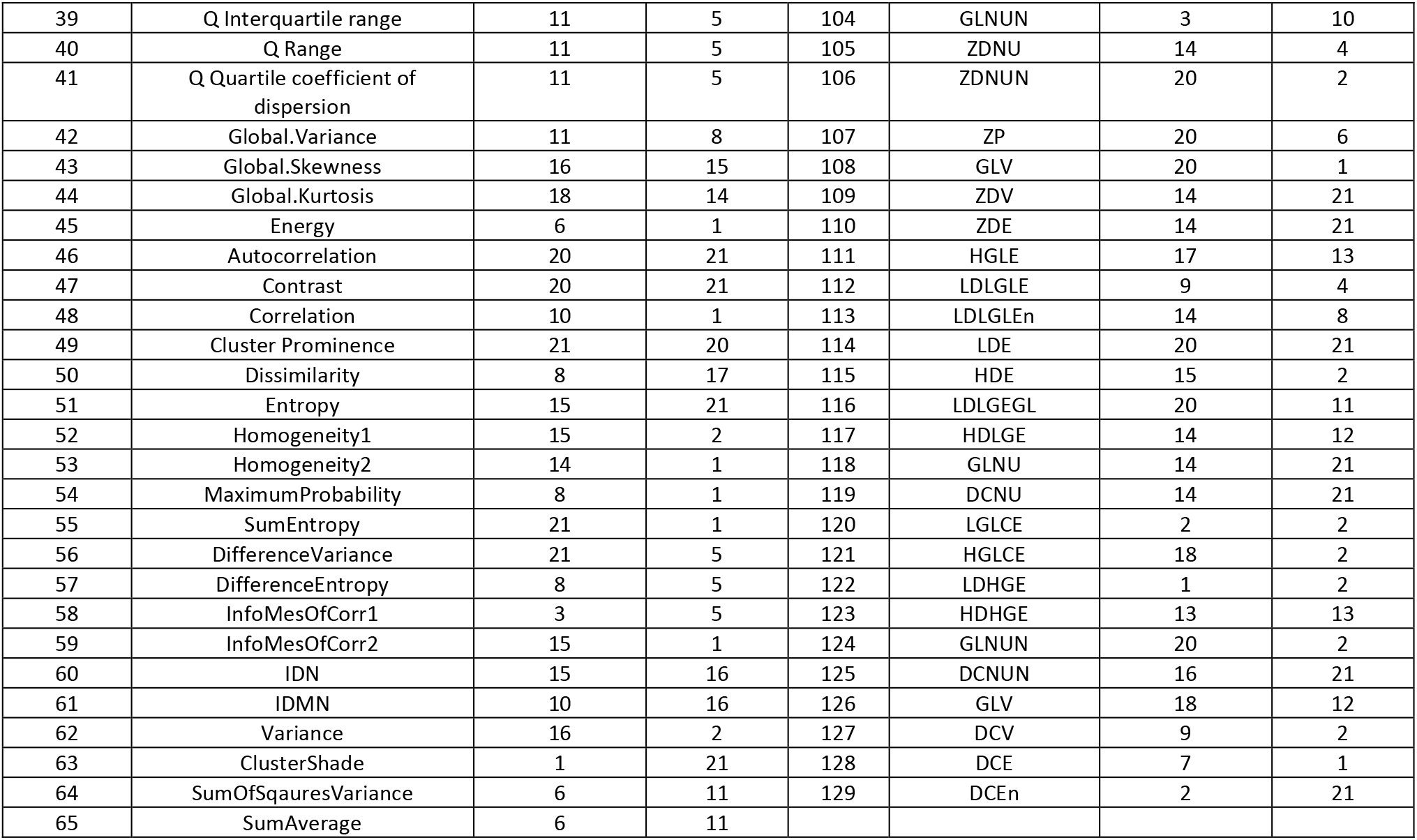
Hierarchical clustering on cross correlation for preclinical and clinical features (dendrogram with distances 3 unit)

**Supplemental Table S3:** AIC and BIC stats for volume dependent radiomic features

| Features name | rho(p) | Stats | Linear | Exponential | Log | Power | Polynomial |
| --- | --- | --- | --- | --- | --- | --- | --- |
| Q Energy | 0.98 | AIC | **459.3** | 577.0 | 531.2 | 461.9 | 460.5 |
|  |  | BIC | **477.6** | 595.3 | 549.6 | 480.3 | 485.0 |
| GLCM.Correlation | -0.98 | AIC | -1281.3 | -1299.1 | -1193.8 | **-1304.3** | -1229.8 |
|  |  | BIC | -1262.9 | -1280.8 | -1175.4 | **-1285.9** | -1205.3 |
| GLCM.Contrast | -0.97 | AIC | -642.2 | -648.3 | -681.5 | **-686.7** | -658.3 |
|  |  | BIC | -623.8 | -629.9 | -663.1 | **-668.3** | -633.8 |
| GLSZM.GLV | -0.98 | AIC | -945.1 | -954.8 | -993.0 | **-905.8** | -811.3 |
|  |  | BIC | -926.7 | -936.4 | -974.7 | **-887.5** | -786.8 |
| GLSZM.ZSV | -0.96 | AIC | -822.1 | -832.7 | -868.6 | **-925.9** | -830.2 |
|  |  | BIC | -803.7 | -814.3 | -850.2 | **-907.5** | -805.7 |
| NGTDM.Busyness | 0.97 | AIC | **-68.5** | -16.4 | -21.3 | -60.0 | -60.8 |
|  |  | BIC | **-86.9** | -34.8 | -39.6 | -78.4 | -85.3 |
| NGTDM.Strength | -0.97 | AIC | -53.6 | -63.2 | -118.8 | **-156.6** | -86.1 |
|  |  | BIC | -35.3 | -44.8 | -100.5 | **-138.2** | -61.6 |
| GLDZM.GLNU | 0.97 | AIC | 129.7 | 218.9 | 171.9 | **121.5** | 137.7 |
|  |  | BIC | 148.0 | 237.3 | 190.3 | **139.9** | 162.2 |
| NGLDM.GLNU | 0.99 | AIC | **340.7** | 772.6 | 728.3 | 477.2 | 334.9 |
|  |  | BIC | **359.1** | 791.0 | 746.7 | 495.6 | 359.4 |
| NGLDM.DCNU | 0.99 | AIC | **772.3** | 993.0 | 958.3 | 1079.6 | 752.6 |
|  |  | BIC | **790.7** | 1011.3 | 976.7 | 1098.0 | 777.1 |

\* Selected model is highlighted in bold

**Supplemental Table S4:**
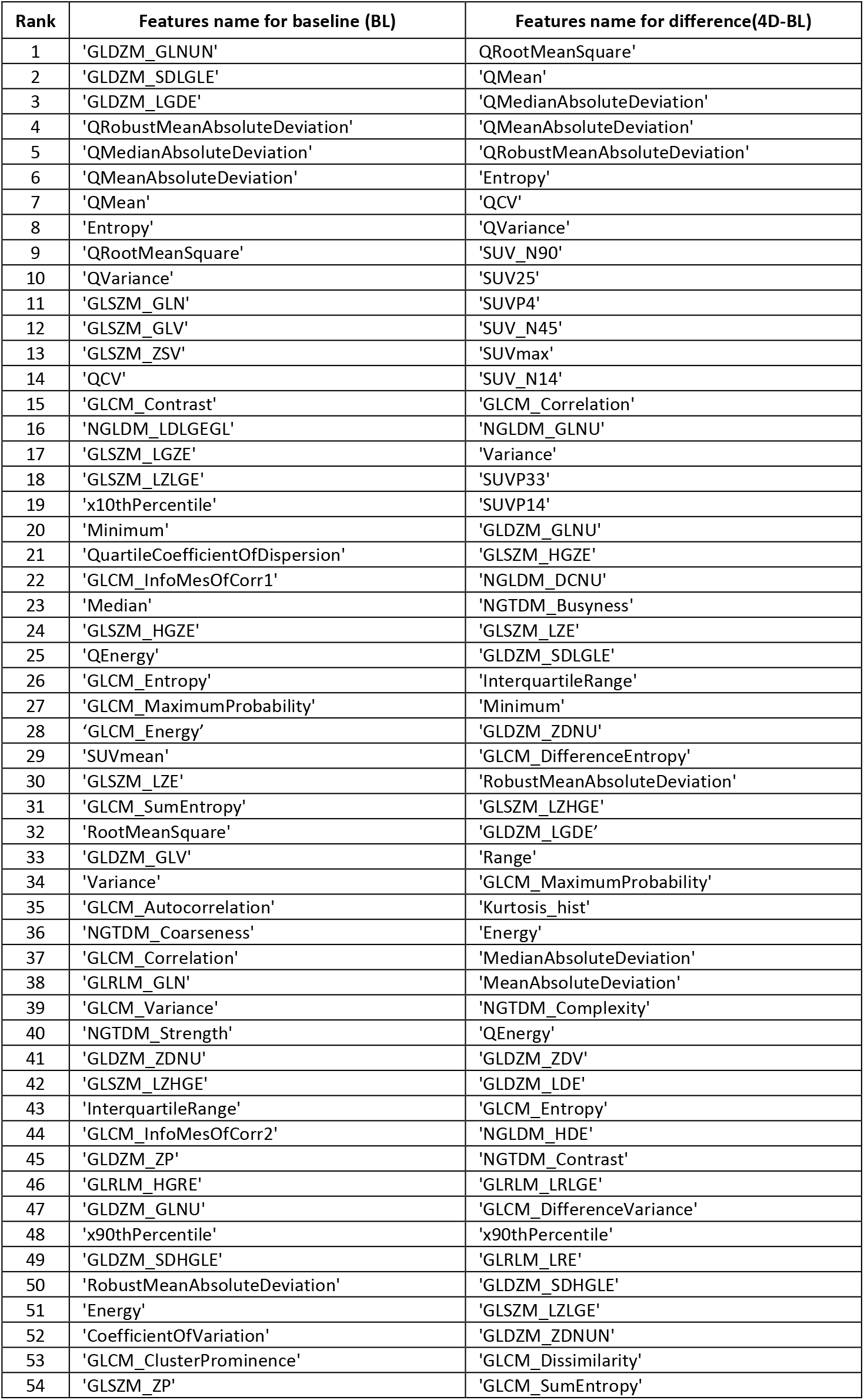
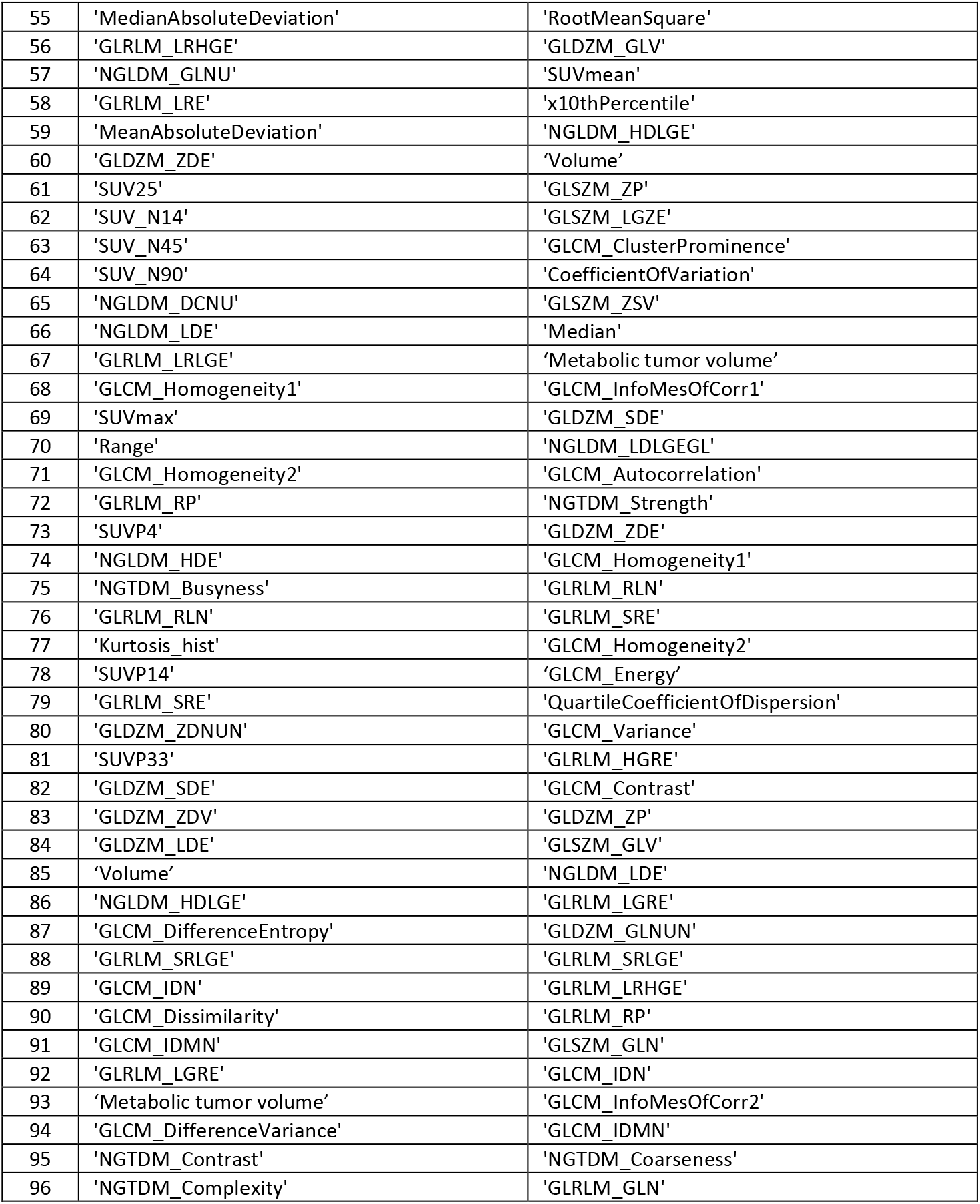
Rank importance (Using ReliefF) of radiomic features based on treatment response prediction (94 reproducible features, tumor volume and metabolic tumor volume)

**Supplemental Table S5:** Accuracy by TNBC Subtyp (%)

| Subtype | CART |  | Naïve Bayes |  | SVM |  |
| --- | --- | --- | --- | --- | --- | --- |
|  | Baseline | Difference | Baseline | Difference | Baseline | Difference |
| IM | 84.37 | 77.75 | 91.25 | 78.5 | 86.5 | 76.25 |
| BL2 | 72.72 | 70.4 | 83.5 | 73.25 | 75.55 | 71.72 |
| M | 81.25 | 71.25 | 81.25 | 82.5 | 83.75 | 75 |
| BL1 | 87.7 | 68.25 | 96.25 | 85.25 | 91.25 | 82.5 |
| LAR | 62.5 | 75.00 | 72.5 | 82.5 | 65 | 75 |

**Supplemental Table S6:** Performance of ML algorithms

|  |  | Preclinical |  |  |  | Clinical |  |  |  |
| --- | --- | --- | --- | --- | --- | --- | --- | --- | --- |
|  |  | Prediction |  | Assessment |  | Prediction |  | Assessment |  |
|  |  | NB | SVM | NB | SVM | NB | SVM | NB | SVM |
| RadSig | F-score | 86.93 | 82.62 | 78.78 | 75.30 | 66.67 | 72.00 | 64.94 | 70.00 |
|  | TPR | 85.81 | 83.33 | 78.00 | 77.38 | 75.00 | 75.00 | 56.82 | 63.64 |
|  | TNR | 86.67 | 78.59 | 78.57 | 72.94 | 53.85 | 69.23 | 80.00 | 80.00 |
|  | PPV | 88.08 | 81.92 | 79.59 | 73.33 | 60.00 | 69.23 | 75.76 | 77.78 |
|  | Accuracy | 86.21 | 81.14 | 78.26 | 75.13 | 64.00 | 72.00 | 67.86 | 71.43 |
| SUV<sub>mean</sub> | F-score | 53.24 | 62.30 | 58.82 | 76.47 | 51.06 | 61.54 | 50.00 | 56.41 |
|  | Sensitivity (TPR) | 52.00 | 63.33 | 76.92 | 86.67 | 50.00 | 66.67 | 45.45 | 50.00 |
|  | Specificity (TNR) | 53.57 | 57.14 | 31.25 | 57.14 | 57.69 | 53.85 | 60.00 | 70.00 |
|  | Precision (PPV) | 54.55 | 61.29 | 47.62 | 68.42 | 52.17 | 57.14 | 55.56 | 64.71 |
|  | Accuracy | 52.76 | 60.34 | 51.72 | 72.41 | 54.00 | 60.00 | 52.38 | 59.52 |
| SUL<sub>peak</sub> | F-score | 48.28 | 64.41 | 78.26 | 76.54 | 52.17 | 55.32 | 53.66 | 55.42 |
|  | Sensitivity (TPR) | 50.00 | 65.52 | 84.38 | 83.22 | 50.00 | 54.17 | 50.00 | 52.27 |
|  | Specificity (TNR) | 46.67 | 62.07 | 61.54 | 63.83 | 61.54 | 61.54 | 60.00 | 60.00 |
|  | Precision (PPV) | 46.67 | 63.33 | 72.97 | 70.86 | 54.55 | 56.52 | 57.89 | 58.97 |
|  | Accuracy | 48.28 | 63.79 | 74.14 | 73.79 | 56 | 58 | 54.76 | 55.95 |
| SUV<sub>max</sub> | F-score | 60.00 | 63.64 | 77.43 | 75.36 | 48.00 | 59.26 | 40.00 | 42.11 |
|  | Sensitivity (TPR) | 60.00 | 63.23 | 86.67 | 86.67 | 50.00 | 66.67 | 36.36 | 36.36 |
|  | Specificity (TNR) | 57.14 | 59.26 | 60.14 | 53.57 | 46.15 | 46.15 | 50.00 | 60.00 |
|  | Precision (PPV) | 60.00 | 64.05 | 69.97 | 66.67 | 46.15 | 53.33 | 44.44 | 50.00 |
|  | Accuracy | 58.62 | 61.38 | 73.86 | 70.69 | 48.00 | 56.00 | 42.86 | 47.62 |

